# Evidence for the Presence of Hyphae and Fruiting Body Calcium Oxalate Crystallites in Schizophyllum commune

**DOI:** 10.1101/2021.08.13.456293

**Authors:** Xiangyue Xiao, Tianji Huang, Jingyi Zhang, Qiuyi Su, Lin Tao, Xiaojie Chen, Yuchen Zhang, Yufan Jiang, Yan Ding, Xiaoming Yu, Wei Liu, Hong Ji, Chi Song

## Abstract

Biomineralization is a phenomenon in which organisms form crystals. Studies have shown that many fungi have the ability to biomineralize, it can exhibit calcium oxalate crystals on their hyphae and fruiting body. *Schizophyllum commune* is a common saprophytic fungus distributed all over the world, but there is little research on its biomineralization. In this paper, *S. commune* fruiting body from three different provinces of China were collected, and isolation for hyphal cultured to obtain several samples. Utilizing light microscope, FE-SEM, and EDAX, the existence of crystals on the fruiting body and mycelium of each strain was found, and their morphological characteristics and ion content were analyzed. It was ultimately established that biomineralization occurs on *S. commune.*

## Introduction

Biomineralization refers to the process by which organisms form minerals. A wide array of biologically induced minerals can be crystallized, a few others might formed amorphous structure, and calcium carbonate minerals are the most abundant biogenic minerals[1]. Among them calcium oxalate(CaO_X_), which is formed when the product of the concentrations of oxalate and calcium ions is greater than the solubility product of calcium oxalate[2] Many plants[3] and some bacteria[4] are capable of producing CaO_X_. Production of CaO_X_ has also been reported in lichenous, mycorrhizal, and litter fungi[5].

Many fungi have ability to exhibit CaO_X_ crystals on their hyphae and fruiting body. The ubiquity of CaO_X_ crystals on fungal hyphae suggests that their formation may provide a selective advantage to the organism[5]. In plant, Various functions have been ascribed to CaO_X_ crystals and these diverse functions depend on crystal amount, distribution, and morphology as well as features of the cells that produce them[3]. For fungi, the function of calcium oxalate has yet to be established, but by speculating on the function of calcium oxalate in plant, CaO_X_ formation is hypothesized to regulate intra-cellular pH and levels of oxalate and Ca and, hence, serves as a major sink for toxic amounts of Ca in soil and other environments[6].

Cystidia are specialized sterile cells found in the Agaricomycetes, Basidiomycota[7]. Cystidia have been thought to function as space makes in the hymenium, as storage cells, and its distribution throughout the various tissues, including the gills or hymenial tissue[8]. Many mushrooms produce hymenial cystidia with apical crystals[9]. But how the crystal formation and the crystal function on cystidia were rarely been illuminated.

*Schizophyllum commune* is a species of fungus in the genus Schizophyllum. which is also called split gill mushroom and “white ginseng” by local people in Yunnan province of China[10], The mushroom resembles undulating waves of tightly packed corals.‘‘Gillies” or Split Gills vary from creamy yellow to pale white in color. The cap is small, 1–4.5 cm wide with a dense yet spongey body texture. It is one of the most common fungi, widely distributed in temperate and tropical regions, and commonly occurs as a weak parasite or saprophyte on a wide range of woody plants and occasionally on herbaceous plants[11]. *S. commune* mainly adopts a saprobic lifestyle by causing white rot[12]. At least 150 genera of woody plants are substrates for *S. commune*, but it also colonizes softwood and grass silage. The mushrooms of *S. commune* that form on these substrates are used as a food source in Africa and Asia[13].

So many researches, including genetics, genome, pharmacology and also cytology, have been carried out look for instance of the genetics of *S. commune*, which reported by John Raper[14]. However, there is no report on the biomineralization of this model fungus.

In order to better understand the structural characteristics of this model organism, we identified the morphological and molecular characteristics of *S. commune* and their strains from Mianyang, Zhumadian, and Wuhan.Explore whether it can form minerals and produce oxalate such as CaO_X_, tested the hypothesis that *S. commune* can be biomineralized.

## Materials and methods

### Strains

*S. commune* fruiting body were collected in different wild areas (Mianyang in Sichuan province; Zhumadian in Henan province, and Wuhan in Hubei province). Experimental strains obtained by conventional sterile tissue separation which were identified by morphological and molecular characteristics. Using a sterile punch, mycelium were inoculated (5 mm) under the aseptic condition and grown in Petri dishes (10*1.5 cm) containing 25 ml of the potato-dextrose agar (PDA)medium, kept the petri dishes covered and placed in biochemistry cultivation cabinet which temperature remained at 25°C for 7 days. All samples were kept stationary during the course of experiment.

When needed, some sterile coverslips were inserted into the plates containing PDA to obtain mycelium cultured from the petri dish. All the tested strains and specimens were deposited in the Spawn Test Center of the Institute of Applied Mycology at the University of Huazhong Agricultural, China.

### Light Microscopy

After the incubation period, the petri dishes were opened and the glass slides were removed and immediately observed under normal and polarized white light with an Olympus BX51 optical microscope. The relevant information was photographed with Kodak ProImage 100 film and the scales were obtained with the projection of a micrometric slide under the same conditions utilized in the illustrations.

### Strain identification

The mycelia of the tested strains of *S. commune* were grown in liquid CYM medium at 25°C for 7 Days. Genomic DNA was extracted from mycelium frozen in liquid nitrogen following CTAB method[15]. DNA concentration and purity were determined with the UV-1700 spectrophotometer (Shimadzu, Japan). Qualified samples were diluted to 50 ng/μl for PCR amplification. Two oligonucleotide fungal primers described by White[16] were used for amplification. The ITS region primers(ITS 1, 5’-TCC GTA GGT GAA CCT GCG G-3’; ITS 4, 5’-TCC TCC GCT TAT TGA TAT G-3’) make use of conserved regions of the 18S (ITS 1), and the 28S (ITS 4) rRNA genes to amplify the intervening 5.8S gene and the ITS 1 and ITS 2 noncoding regions.

The PCR assay was performed with 1μl of test sample in a total reaction volume of 20 μl consisting of PCR buffer; 0.1 mM (each) dNTPs; 1.5 mM MgCl_2_; 0.25 mM (each) primer; and 1.5 U of rTaq DNA polymerase (TaKaRa, Japan). Thirty-five cycles of amplification were performed in a Stratagene Robocycler model 96 thermocycler after initial denaturation of DNA at 95°C for 5 min. Each cycle consisted of a denaturation step at 95°C for 30 s, an annealing step at 58°C for 30 s, and an extension step at 72°C for 1 min, with a final extension at 72°C for 10 min following the last cycle. After amplification, the products were analyzed by electrophoresis in 1%(w/v) agarose/TAE gels, and detected by ethidium bromide staining using GL200 image analysis system (Kodak, USA). The unique DNA fragment was excised from the agarose gels and purified using DNA gel extraction kit (Doupson, China). The purified fragment was sequenced directly using ABI 3730 DNA sequencer. The ITS sequences successfully sequenced were identified by the National Biotechnology Information Center (NCBI)database, determined the experimental strain is the *S. commune* strain.

### Electron microscopy

The mycelium cultured on insert coverslip and some wild mushroom fruiting body section samples were eluted twice with PBS buffer, after two hours, placed it in 4% glutaraldehyde to fixed. Fixed samples dehydrated through graded alcohol (25, 50, 75, 90, 100, 100%) 10-15min every treatment, and 100% ethanol was replaced by tert-butyl alcohol, and then absolutely drying though vacuum freeze-drying. Some samples were dried more than 24h in the oven for comparison the above chemistry method for checking whether the crystal would be destructed during the drying process. Dry samples were gold-coated (10 nm) and observed using a Hitachi S-4800 FE-SEM (Field Emission Scanning Electron Microscope).

### Energy Spectrum Analysis

The energy spectrum of the area to be measured were investigated by an Energy-dispersive X-ray spectrometer (EDAX), operated at 15kV. For each sample, multiple point scanning and surface scanning of different regions of interest was carried out, and the noncrystalline regions were scanned at the same time. Each specific region was scanned three times repeatedly. Fully automated data collection and normalized with the use of the energy spectrum analysis software that comes with the energy spectrum, and the element quantitative analysis was carried out according to the standard data in the database, and all the results were displayed by weight percentage.

### Determination of Ca ions in the fruiting body of wild schizophyllum

Calculate the concentration of calcium in fruit body samples collected in different regions by ICP-MS. First, dry the sample to constant weight. After weighing a certain amount of fruiting body specimens, the samples were digested by microwave digestion. Then, dilute to 5 ml with 2.5% nitric acid. Using high performance liquid chromatography coupled with inductivity coupled plasma mass spectrometry (HPLC-ICP-MS), make a standard curve of calcium ions standard solution, and determine the calcium ion concentration of each sample. All experiments were repeated three times.

## Results

*S. commune* is a wood saprophytic fungus widely distributed all over the world. Year-round occurrence on decaying hardwoods. Fruiting Body diameter 0.6-4.2 cm; fan-shaped when attached to the side of the log; irregular to shell-shaped when attached above or below; upper surface covered with small hairs, dry, white to grayish or tan; under surface composed of gill-like folds that are split down the middle, whitish to grayish; stem short or absent; flesh tough, leathery, pallid. (Fig. 1). Besides, the mycelium has a lock-like joint, the small rod-shaped protrusions are visible on the mycelium. The top of the protrusion is a hollow sphere, the height of the protrusion is 2-4μm, and the diameter of the sphere is 2.5μm (Fig. 2).

**Fig. 1.**
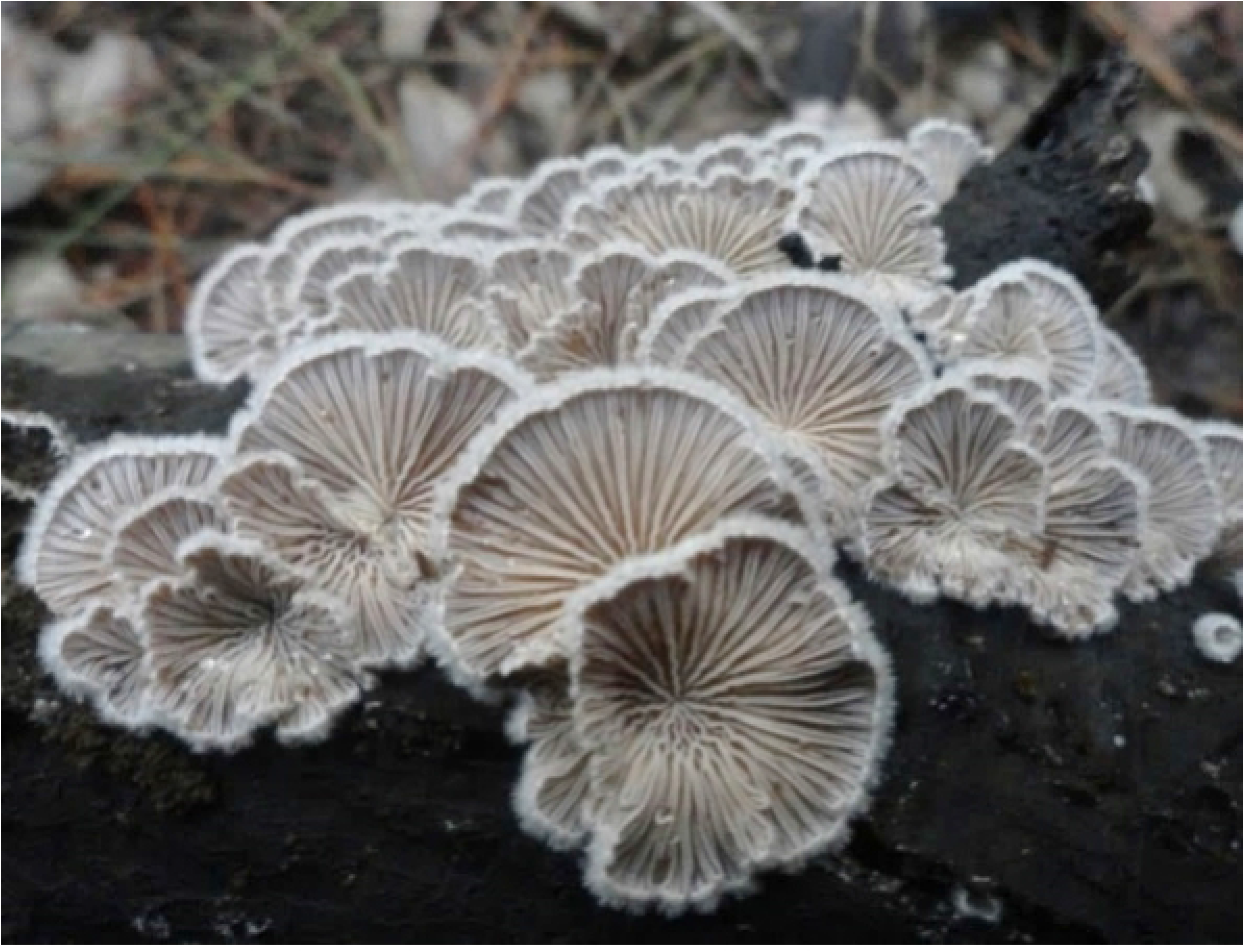
S. commune fruit body

**Fig. 2.**
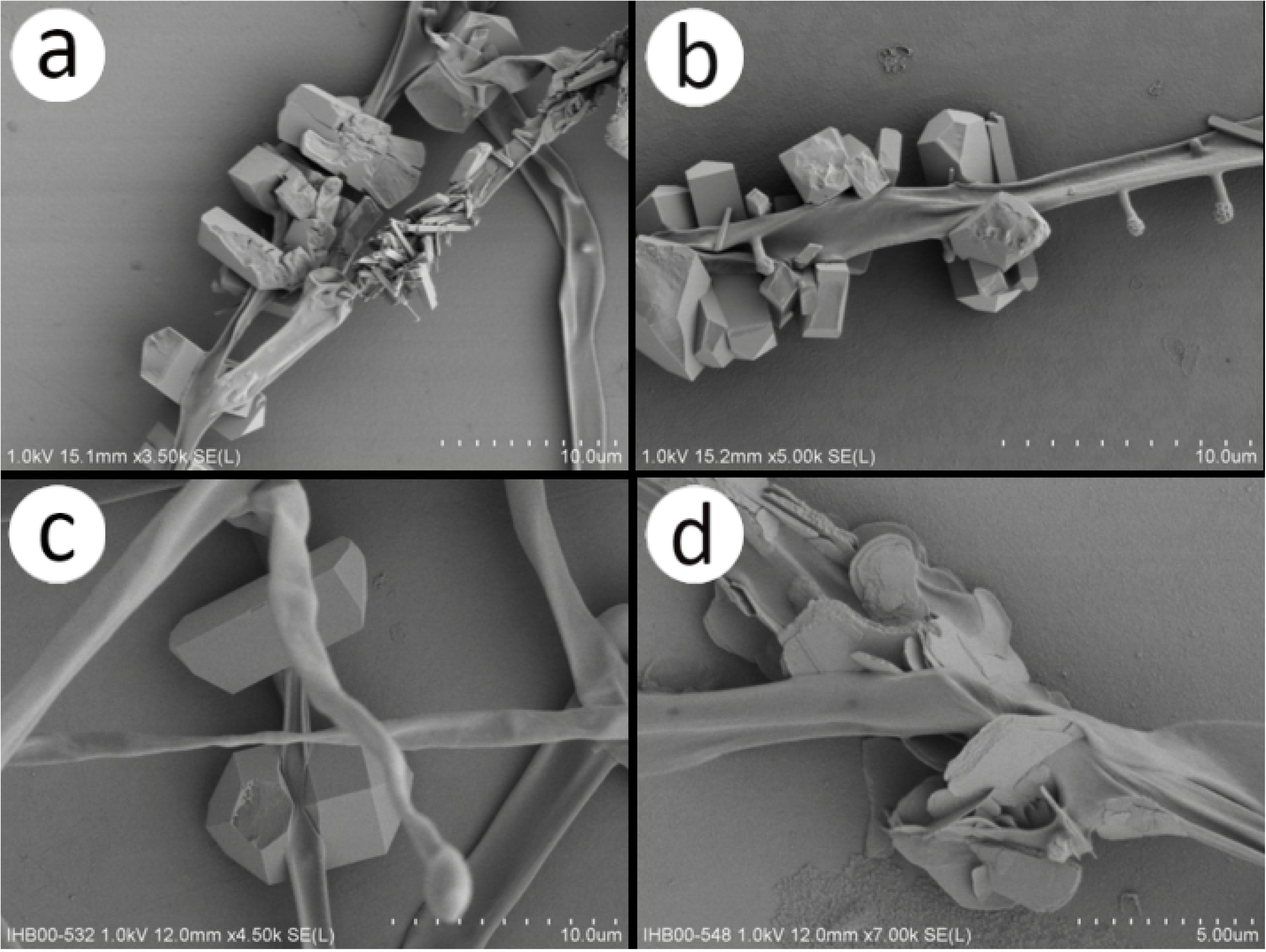
a, b, c, d are photomicrographs of different locations of the mycelium. It can be seen that there are obvious biomineralized crystals on the surface of the mycelium existing in clusters with irregular shapes. The scale bar of a, b, c is 10 μm, the scale bar of d is 5 μm.

ITS sequence analysis displayed by NCBI database alignment, The DNA sequence of the isolated strain is more than 98% similar to the DNA sequence of *S. commune* in the database, there is no more similar option than this sequence. The comparison results showed that the tested strain was a *S. commune* strain.

Observation and analysis of mycelium cultured from a petri dish by optical microscope. There are a large number of clustered substances on the mycelium that are approximately strips, prisms, or cuboids. The DIC differential imaging shows that they have special light-shielding, it was observed to be transparent with transmitted light. Through the scanning electron microscope observation of mycelium, these clusters of long strip objects have different shapes, such as long strip, Bar shaped dodecahedron, irregular flake, and irregular trapezoid. The long crystal has non-uniform ends, the longest is about 8um long, and the width varies from 0.2 to 0.8um; the largest crystal can be 10um long and 7um wide, and some mycelium can be observed to penetrate through the middle of the crystal (Fig. 2).

Observation of *S. commune* fruit body by scanning electron microscopy found that there were a large number of orthocrystals on the basidiomycete sublayer cystoid hyphae of the fruit body, these crystals have various shapes. Most of the small crystals are irregular but non-flaky. The large crystals are mostly octahedral with clear contours, sharp edges, and flat surfaces. There are no crystals on the mycelium of other hymenium (Fig. 3).

**Fig. 3.**
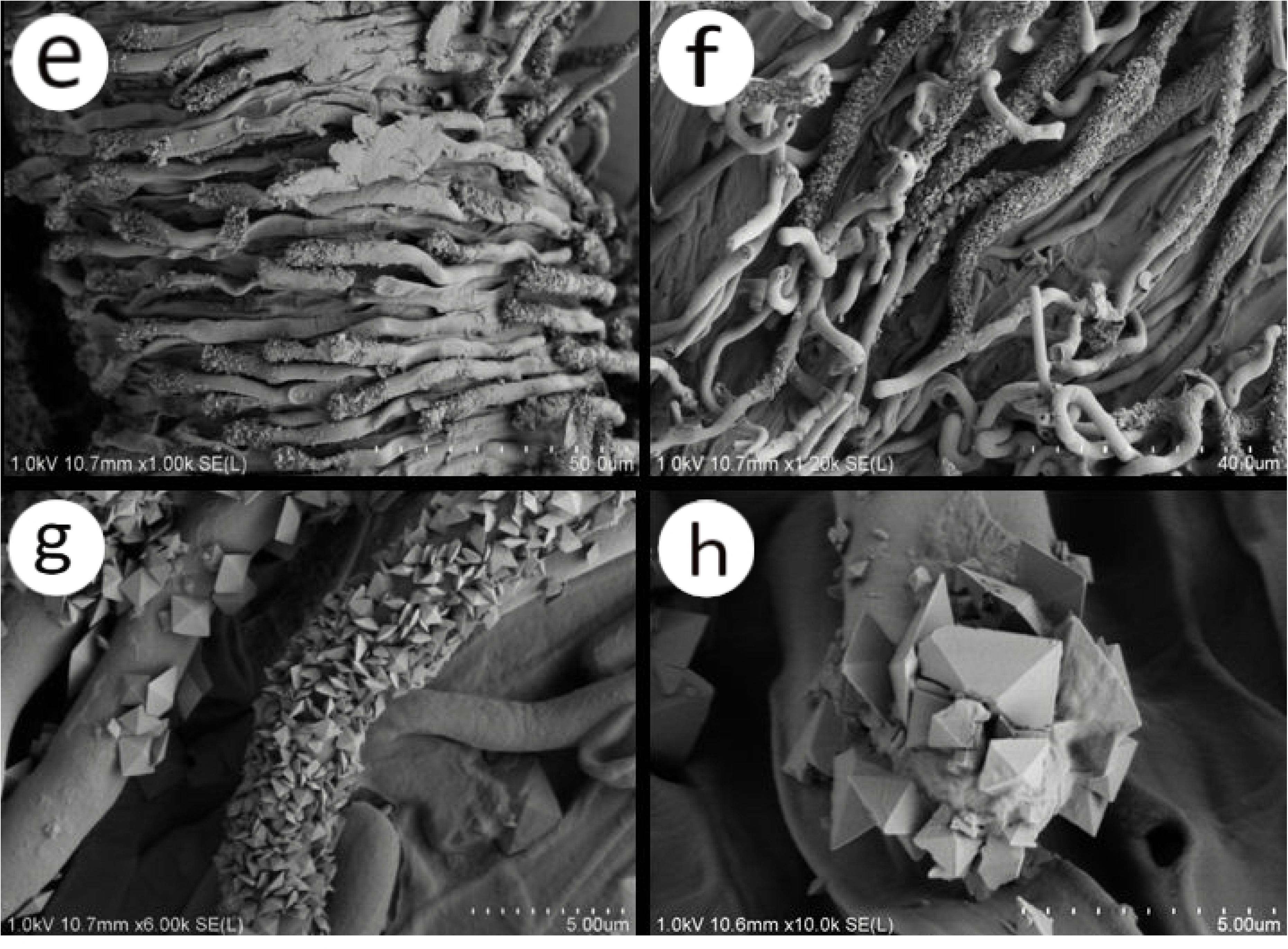
e, f, g, h are micrographs of hyphae at the site of a basidiomycete sublaminar cystidium of the *S. commune* fruiting body. The scale bar of e is 50μm. It can be seen that the crystallization of the top of the mycelium is obvious. The scale bar of f is 40 μm. It can also be seen that the closer to the top of the hypha, the denser the crystal clusters. The scale bar of g and h is 5μm, and the morphological characteristics of crystals on the hypha can be clearly seen.

Under laboratory conditions, Energy spectrum analysis of the crystals of mycelium cultured on the inserted slides showed that the presence of different ions could be detected, namely C, O, Na, Mg, Al, Si, S, Cl, K, Ca, Br, Cr, Zr, Pd, W, and Pt. The detected elements with percentage content greater than 2% are C, O, Si, Ca, W, Pt. Exclude Pt ions introduced by gold spraying(Pt gold and palladium)treatment during the sample preparation process, and Si(SiO2)ions brought in by Carrier slides of mycelium, the percentage concentration of C, O, and Ca ions in the crystals is higher than that in the control area(in the hyphae or in the glass slide), the C ion content is 11.95% and 7.35%, and the O ion content is 43.95 and 32.82%, Ca ion content is 28.82% and 8.84%(Fig. 4).

**Fig. 4.**
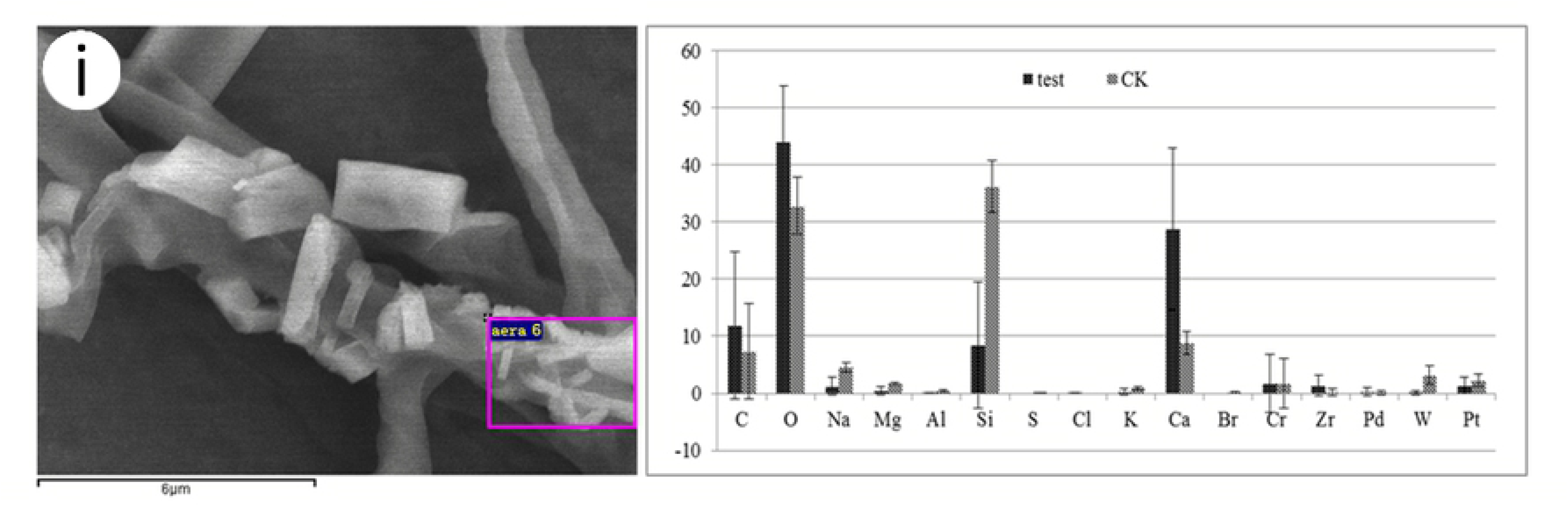
The picture on the left of i is a photomicrograph of a part of the mycelium obtained from the culture of *S. commune*. The scale bar is 6 μm. It can be seen that there are crystals on it. One of the crystals is subjected to energy spectrum analysis. Then select a non-crystalline mycelium site for the same analysis, and finally combine the data to get a comparative histogram of the content of each element(right).

Scanning electron microscope observations of *S. commune* fruit body revealed that there were a large number of crystals on the apical surface of mycelium at the basidiomycete sublaminar cystidium, and no obvious crystallization occurred on other tissue areas including basidiomycetes, basidiomycetes, and skeletal hyphae. EDAX energy spectral scanning analysis using the crystals on *S. commune* fruiting body as experimental group and the *S. commune* tissues as control revealed that the presence of nine ions of C, O, Mg, S, K, Ca, Zr, Pd, and Pt was detected in both the crystals and the tissues. In which the percentage contents of Mg, s, K, and Zr ions in the test and control regions were all less than 2%, excluding Pt ions introduced due to sample preparation, four abundant elements were detected in the crystals of fruiting body, namely C, O, Ca, and Pd. Among them, the content of C ions in the crystal of the experimental group was 0, which was lower than that in the tissue of the control area (the site of data collection or area n = 18) on a 5.39% percentage of C ions; in the experimental group, the percentage of O ions was 49.42%, which was lower than the 60.01% percentage of control; and in the experimental group, the percentage of Ca ions was 35.59%, which was significantly higher than the 13.90% percentage of control (Fig 5).

**Fig. 5.**
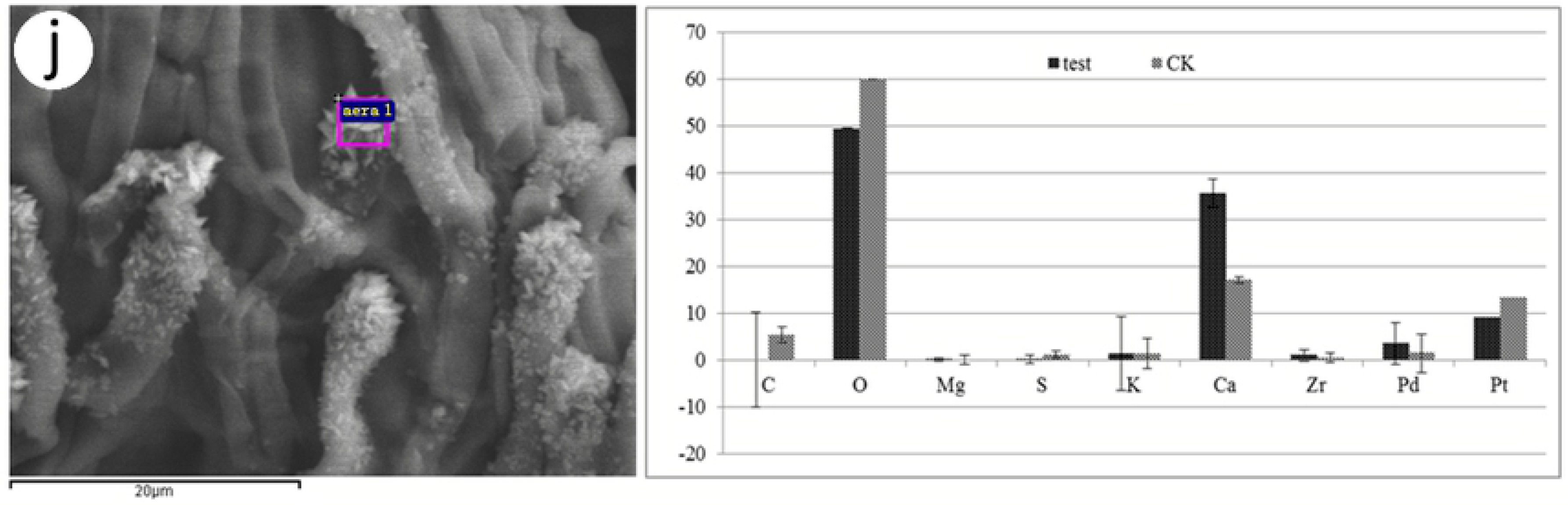
j(left)is a photomicrograph of a part of the fruit body of *S. commune*. The scale bar is 20 μm. It can be seen that there are crystals on it. One of the crystals is subjected to energy spectrum analysis. Then select a non-crystalline fruit body site for the same analysis, and finally combine the data to get a comparative histogram of the content of each element(right).

Scanning electron microscopy results showed that the crystals on the fruiting body mainly existed as octahedra, which may be a special CaO_X_ form, which needs to be further investigated.

Measurements of calcium ion concentrations in wild *S. commune* fruit bodies at three collection sites found that the highest calcium ion content in specimens collected in Wuhan was 322.54 mg/100g (dry material), and the lowest calcium ion content in specimens collected in Zhumadian was 229.85 mg/100g, of which the content of calcium in Mianyang specimens was 236.62 mg/100g. (Table 1)

**Table 1.** Calculate the concentration of calcium in fruit body samples collected in different regions by ICP-MS, and obtain the calcium ion content data of three wild strains.

| Sample | Ca<sup>+</sup> concentration<br>(mg/L) | Sample quality<br>(g) | Actual<br>concentration<br>(mg/g) | Average<br>(mg/100g<br>dry material) |
| --- | --- | --- | --- | --- |
| zhumadian | 33.26 | 0.07 | 2.27 | 229.85 |
|  | 45.35 | 0.10 | 2.34 |  |
|  | 38.23 | 0.08 | 2.29 |  |
| wuhan | 37.45 | 0.06 | 3.33 | 322.54 |
|  | 49.33 | 0.08 | 3.18 |  |
|  | 55.46 | 0.09 | 3.16 |  |
| mianyang | 28.54 | 0.06 | 2.47 | 236.62 |
|  | 53.47 | 0.12 | 2.26 |  |
|  | 86.33 | 0.18 | 2.37 |  |

## Discussion

Regarding the biomineralization of *S. commune*, In the description of *S. commune*, Linder [17] once pointed out that the opposite side of the gill from the hymenium occurred some differently shaped hairs. Occasionally they may be roughened with crystalline deposits, but the deposits are not constant in their occurrence and when present they are not specific[17]. Since Linder did not indicate whether the crystals appeared on the mycelium or the surface of the fruiting body, nor did they describe and identify the crystals in detail. In response to the status quo, we selected *S. commune* strains from different regions and carried out corresponding research.

Correlation analysis results displayed that different shapes of cluster crystals produce on plate culture mycelium and basidiomycete sublayer cystoid hyphae of the fruiting body. performed energy spectrum analysis to crystals on these samples. The analysis data concluded that the calcium oxalate crystals on the mycelium of S. commune are different from those on the fruiting body and contain more variety of elements than those on the fruiting body. Therefore, the relationship between the species of element contained by the crystal and the organization in which it is located awaits further investigation. However, the contents of the three elements C, O, and Ca are both abundant, so it can be determined that these crystals are calcium oxalate crystals.

Crystals such as calcium oxalate are produced presumably to regulate intracellular pH and levels of oxalate and calcium. Yet, the number and morphology of the crystals on different tissues are different, which may originate from different growth status and the degree of differentiation of each tissue [18], Since the three types of samples can only generate crystals on the mycelium cells of basidiomycete sublaminar cystidium and the plate culture mycelium cells, combined with other literature, it is reasonable to speculate that crystallization can only occur on the mycelium cells of *S. commune* in undifferentiated tissues[18]. But overall, the crystal morphology of these three strains from different regions is similar, and their crystal morphology, size and spatial distribution are different from those of other species, such as Prismatic crystals in the bean seed coat, Raphide crystals from a ruptured Pistia raphide idioblast, etc. it can be seen that the formation of crystals is strictly controlled by genetics, and has specificity among various species[19], while a particular species will only form a subset of a certain crystal type or crystal morphology.

From other research reports, we can see that the usual content (mg kg^−1^ dry material) of calcium in wild-growing mushrooms varied from 100 mg kg^−1^ to 500 mg kg^−1^[20], and for some edible mushrooms, such as the species growing wild in Turkey Nesim[1], the content of calcium varied from 189.7 mg kg^−1^ in Polyporus squamosus to 33786.89 mg kg^−1^ in Morchella esculenta species, Than, from our test results, the calcium content of *S. commune* should be varied from 2290 mg kg^−1^ to 3230 mg kg^−1^. Therefore, we can conclude that the calcium content of *S. commune* is significantly higher than some common wild mushrooms. Although it cannot exceed some edible fungi with higher nutritional value, it is still a kind of strain with rich calcium content.

## Conclusion

It was found through experiments that the biomineralization occurred in all three samples of *S. commune* we selected. Use light microscope and scanning electron microscope to observe its plate culture mycelium and basidiomycete sublayer cystoid hyphae of the fruit body, the plate culture mycelium has clusters of cubes or lamellar crystals of various shapes, and the mycelium at the basidiomycete sublaminar cystidium of the fruit body has irregular or octahedral crystals of various sizes. The energy spectrum analysis showed that it was rich in C, O and Ca elements. It can be inferred that the crystals on the mycelium and fruiting body were calcium oxalate crystals. On top of that the Ca content of *S. commune* was generally higher than for other mushrooms.

## Funding and Acknowledgments

This work was supported by the Basic research on agricultural applications in Suzhou city (No. SNG201616).

